# Detecting transcriptional responses and comparing the virulence of *Pseudomonas aeruginosa* cystic fibrosis isolates in a mung bean model

**DOI:** 10.64898/2026.06.06.729815

**Authors:** Sara Franco Ortega, Ezra Herman, Ifigeneia Kyrkou, Helle Krogh Johansen, James W. B. Moir, Clare S. Mahon, Ville-Petri Friman

## Abstract

A mung bean infection model has previously been shown to differentiate between non-virulent and virulent *Pseudomonas aeruginosa* bacteria. However, it remains unclear how plant and bacteria adjust their gene expression during infection and whether the mung bean model can be used to compare the virulence of clinical cystic fibrosis (CF) *P. aeruginosa* lung isolates. Here, we first explored temporal transcriptomics of *P. aeruginosa* PAO1 and mung bean during an infection. We found that bacterial gene expression followed temporal changes, with an increase in the expression of O-antigen biosynthetic genes, chemotaxis, phosphate intake and phenazine production. Mung bean responded by upregulating genes associated with defence mechanisms and downregulating genes involved in the plant development. From the PAO1 perspective, the core-transcriptomic responses in the mung bean were similar to its responses previously observed in wound and excision and in *in vitro* media and sputum models, while differed from those observed in the bronchial cell model. Furthermore, we used the mung bean to assess the virulence of 119 clinical *P. aeruginosa* CF strains originating from the Copenhagen CF clinic. By quantifying bacterial virulence as a reduction in root and shoot growth and weight of the seeds, we found that CF strains isolated at later compared to early stages of lung infections showed higher virulence. This difference corresponded with the higher number of immune modulation-associated virulence genes and lower number of motility and effector genes, present in the genomes of late compared to early isolated CF strains.

**IMPORTANCE:** Our results demonstrate that based on PAO1 transcriptional profile, the mung bean model is similar to *in vitro* and wound infection models but differs from cell and bronchial models. Moreover, the mung bean model can detect virulence differences between clinical *P. aeruginosa* CF strains, making it a potentially useful high-throughput *in vivo* model for bacterial virulence screening.

## INTRODUCTION

*Pseudomonas aeruginosa* is a Gram-negative, opportunistic pathogen and one of the top-listed human pathogens causing hospital-acquired infections (1, 2). *P. aeruginosa* causes acute and chronic infections in immunocompromised patients, including those with burn wounds, sepsis, ventilator-associated pneumonia, cancer, and cystic fibrosis (CF) (3, 4). CF is caused by a mutation in the gene encoding the cystic fibrosis transmembrane regulator protein, a chloride channel which is involved in clearing the mucus (sputum) and bacteria from the airway surfaces (5). When the channel is not working, the mucus becomes dehydrated and accumulates. It induces an excessive inflammatory response and results in a predisposition for bacterial infections, compromising lung function that could lead to increased morbidity and mortality in people with CF (6). *P. aeruginosa* is the major pathogen in CF lungs (7, 8). The infection starts with *P. aeruginosa* strains showing nonmucoid phenotypes where bacteria adopt a planktonic lifestyle (9). *P. aeruginosa,* however easily adapts and evolves to the rapidly changing environment in CF lungs, leading to chronically persistent strains that are more difficult to eradicate or control with antibiotics (10–12). Typical evolutionary changes of chronically infecting *P. aeruginosa* include formation of biofilms, production of adhesins, reduced antimicrobial susceptibility, and changes in lipopolysaccharide (LPS), motility, type 3 secretion system and quorum sensing amongst others (13–20). Moreover, chronic strains are characterised by high phenotypic diversity (21, 22), hypermutability (23) and an increase in antibiotic resistance (24, 25), and they often suffer loss or reduction of acute virulence. While several microbiological screening methods have been developed to quantify variation in *P. aeruginosa* virulence traits (26), quantifying changes in *P. aeruginosa* virulence *in vivo* is still limited by a lack of reliable high-throughput models.

Several infection models have been developed to assess virulence, pathogenesis and/or disease development in acute and chronic *P. aeruginosa* strains. These models include wax moth, fruit fly, nematode, mouse, ferret or pig (27–34). For example, murine models can be used to study acute infections using third-degree burn on the dorsal surface of the mice and subsequent infections, causing rapid sepsis and mortality in 2 days. In contrast, chronic murine models require dorsal excision, infection with the pathogen and covering with adhesive dressing, which ensure wound healing, and delaying the infection which can persist for weeks (35–37). In addition to insects and mammals, *P. aeruginosa* can also infect amoebae and plants (38–40). Plant infections by *P. aeruginosa* have been less studied, but this pathogen has previously been detected in Nicotiana (41), ginseng (42), soybean (43), barley ((44) and in *Arabidopsis thaliana* (45). When infecting plants, *P. aeruginosa* first attaches to the leaf or other surfaces, thereafter invading and colonising the inside of plant through stomata or wounds. This results in disruption of plant cell walls and membranes, causing a systemic infection and ultimately the death of the plant (45). Previously, Garge et al. (46) confirmed that a mung bean model can be used to distinguish between virulent *P. aeruginosa* PAO1 wild-type from a non-virulent quorum-sensing mutant. However, it is not yet clear how bacterial and plant gene expression changes during mung bean infection, and how well PAO1 transcriptional changes reflect its virulence gene expression in the mung bean compared to other virulence models. Moreover, it remains unclear if a mung bean model can differentiate the virulence of acute and chronic *P. aeruginosa* CF isolates.

By expanding the previous work by Garge et al (2018), we explored *P. aeruginosa* PAO1 and mung bean gene expression changes during the infection relative to other virulence models, which included *in vitro* growth media, lung cells, sputum and skin models using burns and dorsal excisions. We focused on PAO1 strain, due to its high virulence in mung bean, having the biggest similarities with CF strains (47), and due to the wide repertoire of publicly available PAO1 transcriptomic datasets, which allowed comparing its virulence with other infection models in terms of gene expression differences. We also assessed whether the mung bean model can be used to distinguish virulence differences between clinical *P. aeruginosa* CF isolates originating from early or late stages of lung infections (12). We found that upon infection, PAO1 increased the expression of several virulence genes including changes expression of O-antigen biosynthetic genes, chemotaxis, phosphate intake and phenazine production. In response, the mung bean upregulated genes associated with disease-related defences and downregulated those genes associated with the growth of the plant. Overall, *P. aeruginosa* transcriptional responses during mung bean infection were similar to *in vitro* and skin infection models but differed from cell and bronchial infection models. Despite this, the mung bean model could differentiate *P. aeruginosa* CF strain virulence based on their virulence gene content and the duration they had been infecting CF patients’ airways, which reflects their evolutionary differentiation during within-patient adaptation. Together, these findings suggest that the mung bean could offer a simple but high-throughput model system to screen the virulence of different *P. aeruginosa* isolates.

## RESULTS

### Virulence-related genes of PAO1 are activated during early stages of mung bean infection

To shed light on the mechanisms that cause infection in mung beans, we first focused on assessing the PAO1 gene expression during early stages (up to 72h) of mung bean infection. Following inoculation with PAO1, we extracted RNA to assess the abundance of viable *P. aeruginosa* in seeds throughout the infection by RT-qPCR. We observed an increase in PAO1 RNA from 24h to 72 h (20.1 pg at 24h, 23 pg at 30h, 75 pg at 48h and 137.2 pg at 72h (Table S8), confirming that bacterial densities were increasing during seed infection. The mapping of RNA reads to the genome assembly of PAO1 produced by this study showed a similar pattern, with 0.59% of the reads belonging to PAO1 at 24h, 0.93% at 30h, 26.76% at 48h and 43.02% at 72h (Table S9). PAO1 quantification with RT-qPCR and by read mapping had a significantly high Pearson correlation (R=0.76, p=0.0065; Figure 1A), confirming that PAO1 was rapidly multiplying within mung bean seeds after the initial infection. Plants also showed disease symptoms, such as growth reduction, change of colour from green to brown and the seeds showed bacterial mucilaginous growth on the surface.

**Figure 1.**
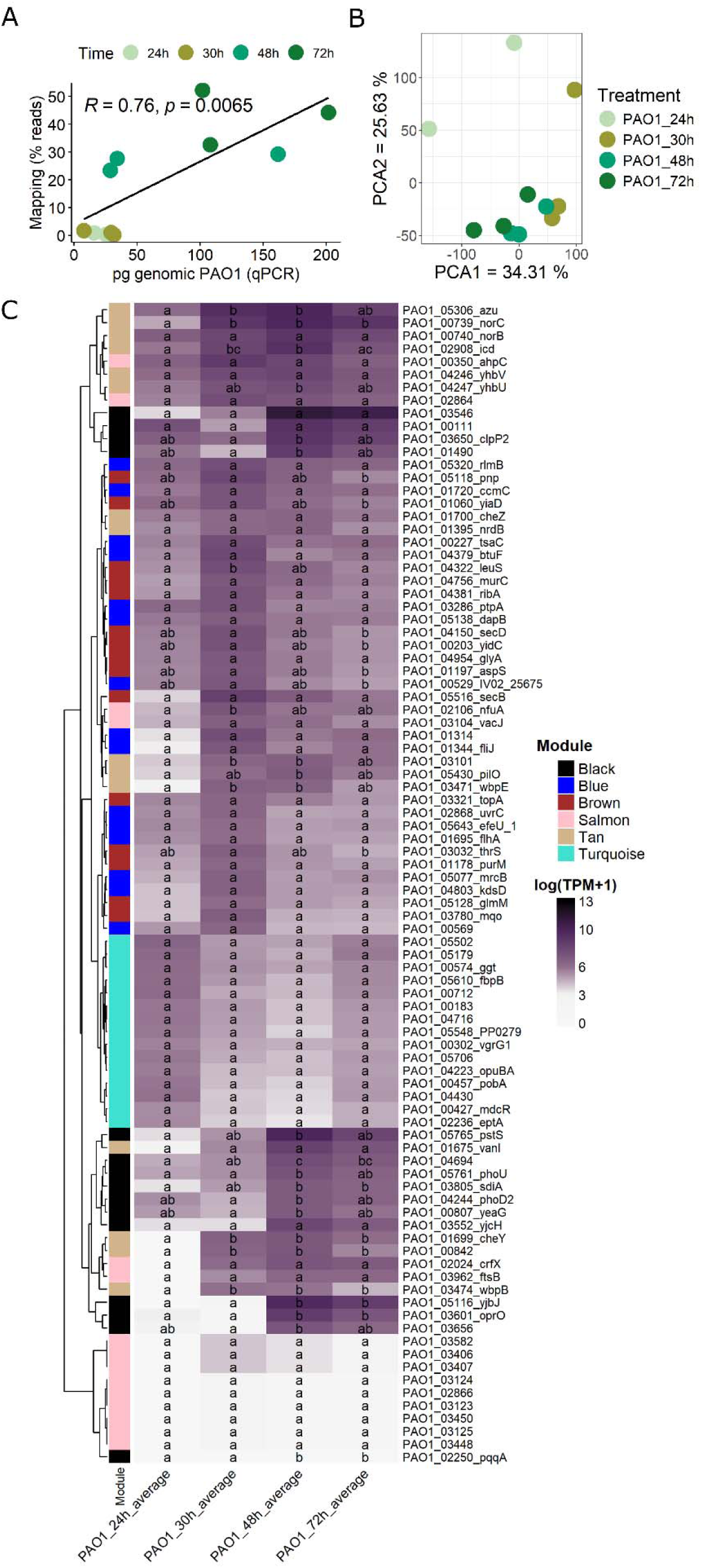
Transcriptional responses of PAO1 during the mung bean infection. **A**. Pearson correlation between the metatranscriptomic read mapping against PAO1 reference genome and the PAO1 qPCR abundances, which confirm the presence of PAO1 at each time point. **B**. PCA plot representing the global PAO1 gene expression changes at the different time points (24 hpi, 30 hpi, 48 hpi, 72 hpi). **C**. Heatmap showing the PAO1 gene expression levels in log scale (transcripts per million (TPMs)+1) focusing on the top 15 genes with the highest connectivity within 6 selected gene coexpression modules associated with PAO1 virulence. These genes included *wbpE* (PAO1_03471), involved in biosynthesis of the central carbohydrate of the B-band O-antigen; *cheY* (PAO1_01699) involved in chemotaxis; *phoU* (PAO1_05761), part of the phosphate regulon; *pstS* (PAO1_05765) involved in phosphate import; *oprO* (PAO1_03601), Phosphate-selective porin; PhzA/B-like protein (PAO1_03656) involved in phenazine biosynthesis; *clpP2* protease (PAO1_03650). Different letters within each cell denote statistically significant differences between time points for each gene, according to a post-hoc Tukey test conducted following ANOVA.

Global changes in PAO1 gene expression over the early stages of the infection were examined using a PCA. The results showed a separation between early versus later stages of infection (24hpi compared to 30hpi, 48hpi and 72hpi), and this separation was especially clear along with the PCA2, which explained 25.6% of the total variance (Figure 1B). Gene expression changes were further analysed using a weighted gene co-expression network (Table S10, Figure 1C), which allowed us to identify 15 gene modules, or clusters of interconnected genes that exhibit highly similar expression patterns across samples. We then performed a gene ontology enrichment analysis to identify gene co-expression modules that were linked with virulence based on enriched GO terms (e.g., functions that help PAO1 adapt to the host, acquire nutrients or invade mung bean immune responses). We found that the black module was enriched for biological processes involved in interspecies interactions between organisms (GO:0044419) and for GO terms associated with cellular responses to starvation (GO:0009267). Genes in the black module could have potentially helped PAO1 interact with the host and survive nutrient-limited conditions inside the mung bean seeds. The blue module was enriched in biological processes involved in the regulation of response to stimulus (GO:004858) and cellular response to oxidative stress (GO:0034599). These genes could hence be associated with life in the host environment and responses to host-induced stress, supporting PAO1 survival and pathogenicity. The brown module was enriched in biological processes involved in symbiotic interaction (GO:0044403) and Gram-negative-bacterium-type cell outer membrane assembly (GO:0043165), suggesting that these genes were potentially associated with pathogen-host interactions and bacterial virulence. The salmon module was enriched with genes linked with cellular response to starvation (GO:0009267), survival in nutrient-limited environments and responses to oxidative stress (GO:0006979), potentially linked with PAO1 responses to plant defences. Finally, the tan module included genes enriched in biological processes involved in cell motility (GO:0048870), while the turquoise module included genes associated with responses to iron (GO:0010039) and peptidoglycan metabolic processes (GO:0000270), which could have been important for pathogen growth and virulence factor expression and modification of peptidoglycan to evade mung bean immune recognition (Table S11).

We next focused on 6 modules potentially associated with PAO1 virulence (black, blue, brown, salmon, tan and turquoise) and identified “hub genes” that had a higher number of connections with other genes within the modules, indicative of their potentially important regulatory role. This was achieved by calculating the number of connections between genes in the module (degree) and identifying genes with high connectivity as hub genes within each module (Table S12-S17). We analysed the gene expression trends and dynamics of hub genes over time to identify transient and sustained gene expression changes. Considering the top 15 highly connected hub genes of the 6 selected modules, we identified *wbpE* (PAO1_03471), which was induced transiently at 30 and 48 hpi (F(_3,7_)=10.4, p-value=0.006; Figure 1C). This gene is involved in the biosynthesis of the central carbohydrate of the *P. aeruginosa* PAO1 B-band O-antigen biosynthesis (48, 49), which acts as a key virulence factor involved in host colonisation (50). We also observed concomitant increase in the expression of *cheY* (PAO1_01699) (F(_3,7_)=124.1, p-value=1.96E-06; Figure 1C), which is a chemotaxis response regulator protein that binds to the FliM protein of the flagellar motors when phosphorylated, causing it to rotate (51). We also identified three genes associated with phosphate uptake: *phoU* (PAO1_05761), part of the phosphate regulon; *pstS* (PAO1_05765), which forms part of the ABC transporter complex PstSACB involved in phosphate import and the porin oprO (PAO1_03601), which is expressed under phosphate-starvation conditions in *P. aeruginosa* (52). Importantly, in *P. aeruginosa*, phosphate starvation causes transcriptional changes that shift to a virulent phenotype (53, 54), indicating that this shift took place at 48hpi (F(_3,7_)=7.2, p-value=0.015; F(_3,7_)=5.9, p-value=0.025; F(_3,7_)=21.3, p-value=6.8E-04 for PAO1_05761, PAO1_05765 and PAO1_03601, respectively; Figure 1C). We also identified increased expression of PhzA/B-like protein (PAO1_03656) involved in phenazine biosynthesis at 48hpi (F(_3,7_)=6.4, p-value=0.02; Figure 1C), which is a clinically important virulence factor regulated by quorum sensing (55): phenazines, such as pyocyanin, mediate tissue damage and necrosis in CF lung (56). Finally, a *clpP2* protease (PAO1_03650), which regulates physiology and is involved in regulating alginate production, biofilm formation (57, 58), showed increased expression at 48hpi and 72hpi (F(_3,7_)=7.9, p-value=0.01; Figure 1C). Together, these results demonstrate the activation of several PAO1 key virulence-related genes during early stages of the mung bean infection, including in O-antigen biosynthesis and chemotaxis, phosphate starvation regulation, biofilm formation and phenazine and protease production.

### Immune-related mung bean genes are activated during early stages of PAO1 infection

We also assessed temporal changes in mung bean gene expression in the presence versus absence of PAO1. In the absence of PAO1, mung bean gene expression changed consistently over time along with both PC1 and PC2 (Figure 2A), while no clear temporal trends in gene expression were observed in the presence of PAO1, which suggests that normal plant developmental processes were likely disrupted due to infection. Moreover, we found a high number of upregulated and downregulated genes at the different time points with 825, 878 and 1217 genes upregulated across time points and 1117, 1453 and 2163 genes downregulated at 30hpi, 48hpi, and 72hpi respectively (Table S18-S20).

**Figure 2.**
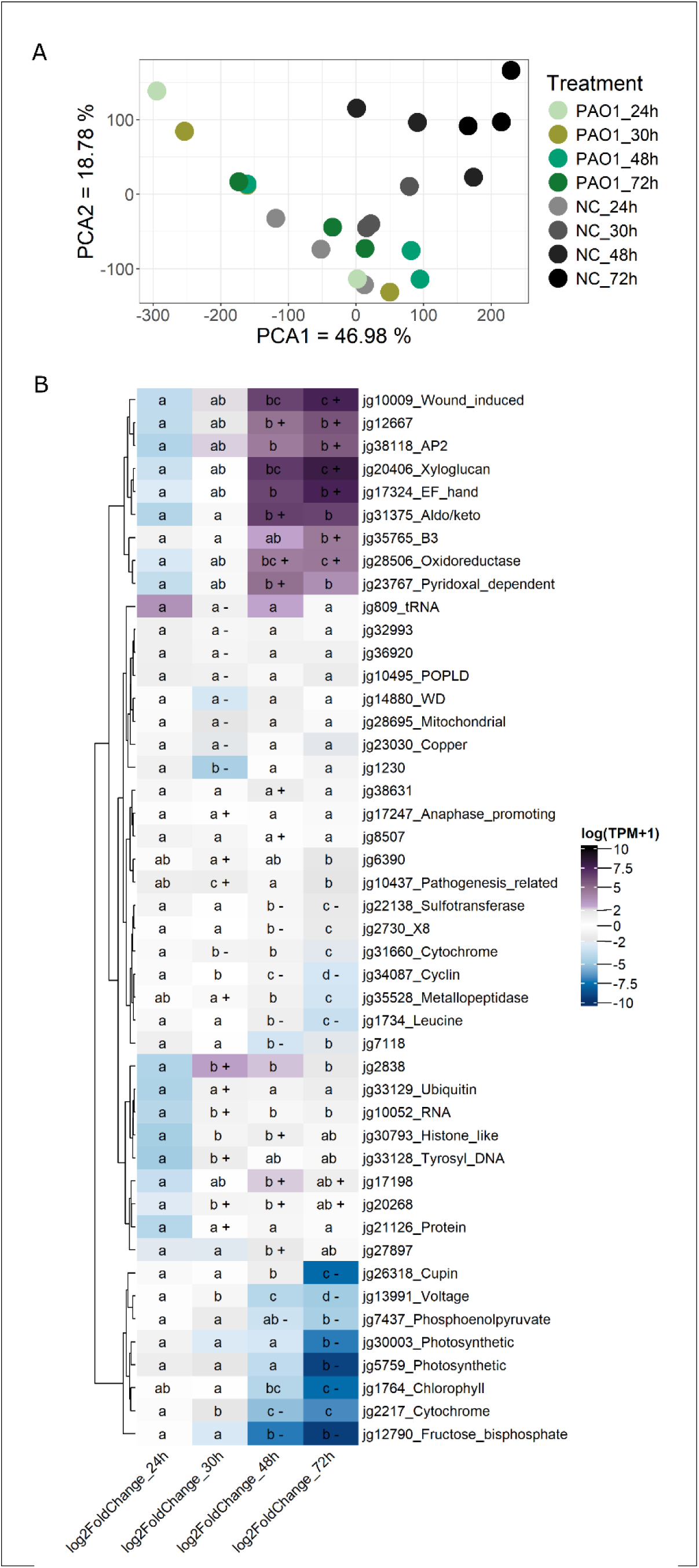
Transcriptional responses of mung bean during PAO1 infection. **A**. PCA plot representing the global gene expression (TPMs) changes in mung bean at different time points (24 hpi, 30 hpi, 48 hpi, 72 hpi) in the presence and absence of PAO1 and negative control (NC). **B**. Heatmap showing the 5 genes with the highest and lowest log2fold changes and the lowest p-adj values across 30hpi, 48pi, and 72hpi time points, including ethylene-responsive transcription factor (jg38118), pathogenesis-related (jg10437), wound-induced protein (jg10009) and B-like cyclin (jg34087). Different letters within each cell of the heatmap denote statistically significant differences between time points for each gene, according to a post-hoc Tukey test conducted following ANOVA. P-values were adjusted with Bonferroni correction. +=indicate log2FoldChange ≥ 2 and padj <= 0.05 and (-) indicates downregulated genes with log2FoldChange ≤ - 2 and padj <= 0.05 at each specific time.

Upregulated mung bean genes in response to PAO1 were enriched with GO terms linked to response to toxic substance (GO:0009636), response to hydrogen peroxide (GO:0042542), response to heat (GO:0009408) and response to wounding (GO:0009611) (Table S21). Within the DEGs, we selected 5 genes with the highest and lowest log2fold changes and the lowest p-adj values across 30hpi, 48pi, and 72hpi time points, resulting in a total of 46 genes. We found an ethylene-responsive transcription factor (jg38118) was upregulated at 72hpi (F(_3,7_)=9.1, p-value=0.008; Figure 2B), indicating this phytohormone are involved in the response to PAO1. Ethylene is a common stress-response hormone often activated in response to pathogens, and for example, in *Arabidopsis*, ethylene biosynthesis is increased in response to the bacterial pathogen *Pseudomonas syringae* pv. Tomato (59). We also found that pathogenesis-related jg10437 gene was transiently upregulated at 30hpi but downregulated at 72hpi F(_3,7_)=32.85, p-value=0.0002; Figure 2B). Finally, we also found a wound-induced jb10009 gene peaking at 72hpi F(_3,7_)=14.91, p-value=0.002; Figure 2B).

The downregulated mung bean genes were enriched in functions such as light harvesting in photosystem I (GO:0009768), chloroplast organization (GO:0009658), chlorophyll biosynthetic process (GO:0015995) and response to light stimulus (GO:0009416). When looking at the most downregulated genes, we identified B-like cyclin gene (jg34087; F(_3,7_)=6371, p-value=2.19e-12; Figure 2B), which is involved in G2/mitotic transition. The downregulation of this gene could be linked to the induction of ethylene-responses, as a reduction in the expression of cycline has been reported in *Arabidopsis* when treating the plants with jasmonic acid (60), a phytohormone that usually acts synergistically with ethylene during defense responses (61).

Together, these findings demonstrate that mung bean upregulated stress and immune-related genes in response to PAO1 infection, indicative of defensive response. Concomitantly, we observed down regulation of genes associated with photosynthesis, which could potentially be explained by a resource allocation trade-off where plant diverted resources from the growth to immune defences (62).

### Comparative transcriptomics identifies similarities between the mung bean and other virulence models

To explore the representativeness of the mung bean plant model, we next compared PAO1 gene expression in mung bean with PAO1 gene expression with other *in vitro* and *in vivo* virulence models. To achieve this, we used publicly available PAO1 transcriptomics data, which was divided into different submodels: media (PIA, M62, MOPs, SCFM, Artificial sputum), in ex-vivo sputum, human bronchial cells, mouse lungs, murine burn wounds, murine chronic wounds and rabbit wound treatments (63–66). After mapping transcripts against the PAO1 genome previously assembled in this study (Table S2 for mapping and Table S22 for TPMs), we divided the gene expression into a core-transcriptome, which included genes expressed at least an average of 1 TPMs in each sample (see material and methods). The treatments were further divided into six different virulence models: *in vitro* media, media+lung cells, sputum, lungs, skin and mung-beans.

When first comparing the expression of all the 5765 PAO1 genes (quantile normalised log(TPMs+1)), we observed that murine chronic and burn wounds and MOPS media showed differential gene expression from all other submodels [60] along the PCA1 axis (Figure 3A, explaining 45.97% of the variance). Moreover, PAO1 responses in the mung bean were statistically different from all the other models along the PCA2 axis (explaining 13.13 % of the variance; perMANOVA, R2=0.15; p-adj=0.015; R2=0.133; p-adj=0.015; R2=0.1; p-adj=0.015; R2=0.11; p-adj=0.01; R2=0.15; p-adj=0.015 for *in vitro* media vs mung bean; media and lung cells vs mung bean; lungs vs mung bean; sputum vs mung bean and skin vs mung bean, respectively).

**Figure 3.**
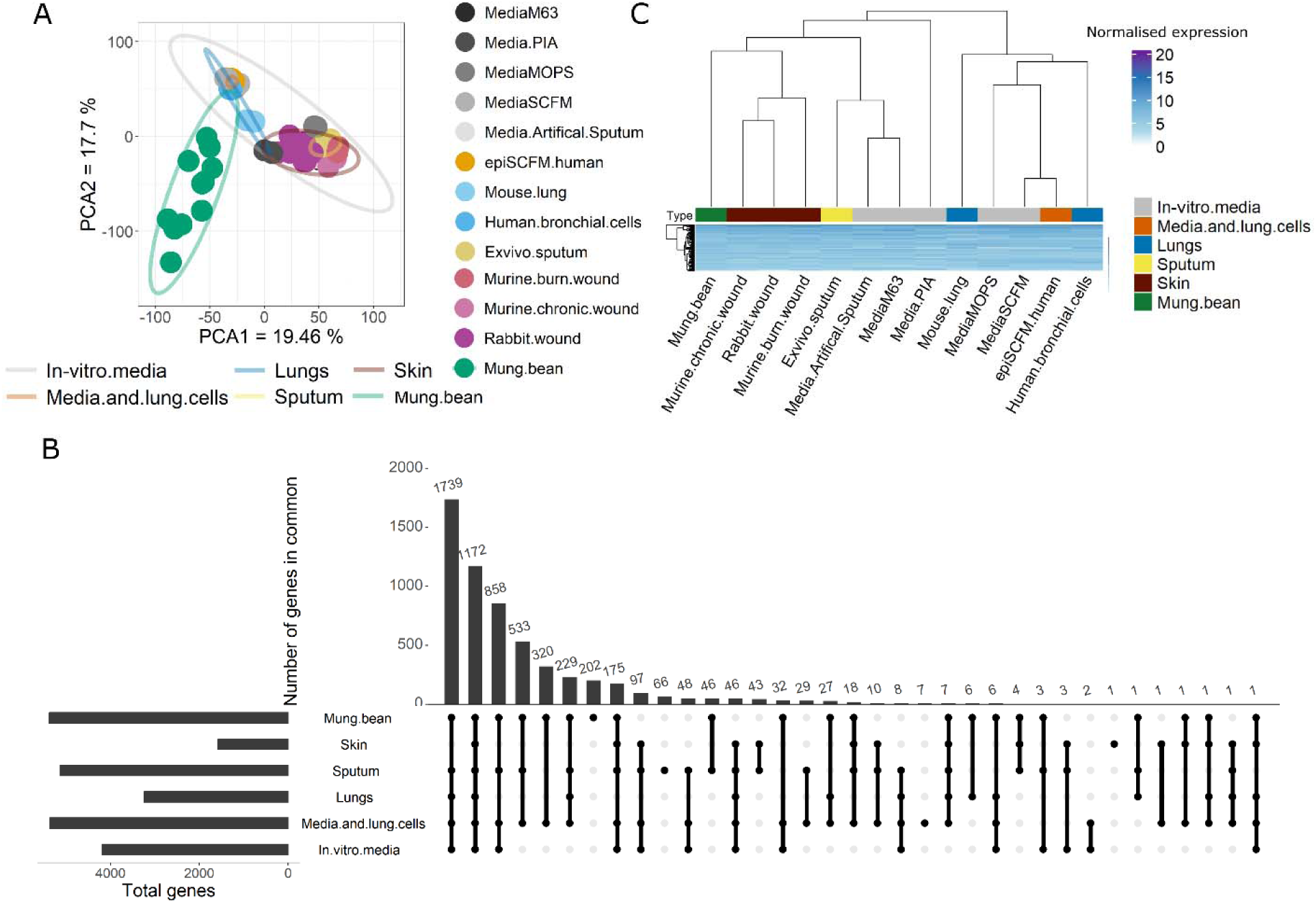
Comparison of the expression of PAO1 genes in mung bean and other virulence models. **A.** PCA of the expression of the quantile-normalised log(TPM+1) 5765 PAO1 genes in the 24 samples (multiple replicates in each; total of 62 samples) belonging to 13 submodels (artificial sputum media, PIA, M63, MOPS, SCFM, epiSCFM. human, human bronchial cells, mouse lung, ex-vivo sputum, murine burn wounds, murine chronic wounds, rabbit wounds and mung beans) and grouped into 6 virulence type of models (*in vitro* media, media+lung cells, sputum, lungs, skin and mung-beans). **B**. Diagram represents the number of common genes shared between models. Core genes were identified as expressed genes across all 6 virulence models considering expression of at least 1 TPM (as explained in material and methods before normalising). **C**. Heatmap of the quantile-normalised log(TPM+1) expression of 1172 core genes expressed in all the 6 models. Columns clustering of the 13 submodels indicate distance between the expression of these genes in the submodel, as a proxy of similarity in expression. Each virulence model is colour-coded in the “type” row.

We then focused on identifying the PAO1 core transcriptome, which consisted of 1172 genes that were all expressed in all the 6 different virulence models (Figure 3B). These genes were enriched in functions such as translation (GO:000641), ribosomal small subunit assembly (GO:0000028), negative regulation of catalytic activity (GO:0043086), and also responses to heat (GO:0009408), SOS response (GO:0009432) and response to osmotic stress (GO:0006970) (Table S23). We then calculated pairwise Pearson correlations of these 1172 genes using quantile-normalised log (TPM+1). The Pearson correlations were then transformed into distance matrix, which allowed us to cluster the gene expression similarities between 13 submodels. These analyses grouped mung beans in a subclade together with murine chronic wounds, rabbit wound and murine burn wounds, which all clustered within a bigger clade containing ex-vivo sputum, artificial sputum media, media M63 and PIA media. However, PAO1 core gene expression in mung bean was clearly different to gene expression observed in human bronchial cells, epiSCFM with human cells, SCFM, MOPS media and mouse lung model (Figure 3C).

Interestingly, we also identified 202 PAO1 genes which were only expressed in the mung bean model. These 202 genes were enriched in response to iron ion (GO:0010039), response to salt (GO:1902074), microcin transport (GO:0042884) and other amino acid transport functions (GO:0006865, GO:0006857, GO:0035672, GO:0015801) (Table S23). At the gene level, these findings were explained by the expression of an operon consisting of *arnD, arnE, arnF* and *arnT* genes, which changes the charge of the lipid A molecule on the LPS (67). Other unique genes included genes of the Npp system, which is homologous to the Yej system proteins in *E. coli* and are involved in the transport and uptake of Microcin C (68, 69) (PAO1_02325 or *yejA* and PAO1_02327 or *yejE*). In Salmonella, Yej operon is upregulated following infection of epithelial cells, which is suggested to be due to the presence of antimicrobial peptides of the innate immune response (70, 71). Consequently, this suggests these PAO1 genes might have been induced in mung bean to counteract plant antimicrobial peptides (72). Moreover, we identified expression of 4 genes belonging to the Type II secretion system (GspEHIK) and induction of phosphatase activity due to the presence of *phoA*. Type II secretion systems could be potentially associated with expression of plant cell wall degrading enzymes as observed with plant pathogenic bacteria (73), while expression of phoA could indicate PAO1 response to low phosphate.

Together these results suggest that PAO1 response in the mung bean model shared some similarities with ex-vivo sputum, *in vitro* media and skin models virulence models, but was less similar to lung cell virulence models. We have also identified 202 genes only expressed in mung beans that can help PAO1 combat plant defenses, for example changing the charge of the lipid A molecule on the LPS, inducting systems that respond to plant antimicrobial peptides and low phosphate and induce the Type II secretion systems potentially to produce wall-degrading enzymes.

### Identifying *P. aeruginosa* genes associated with clinical CF strains’ virulence in mung bean

Previously, Garge et al. 2018 (46) confirmed that infections in mung beans can distinguish wild-type *P. aeruginosa* PAO1 from a quorum-sensing mutant, suggesting this model can detect differences in *P. aeruginosa* virulence. We further explored whether clinical isolates show virulence differences in mung beans and which virulence genes might be associated with the virulence difference observed in mung bean. We used a subset of strains from a CF collection collected in a longitudinal study (12) and compared the virulence of 119 strains in the mung bean model, where the virulence of each strain was tested using 9 seed replicates. The virulence was quantified by measuring bacterial effects on mung bean root and shoot length, seed germination rates (0 if the seeds did not germinate or 1 if the seeds germinated) and weight of the seeds after 5 days post infection (dpi) (higher weight at 5dpi implied lower virulence) (Table S24). Negative control seeds were kept in media without bacterial inoculum [46] and we included PAO1 as a positive control representing a virulent strain (46). PAO1 was genetically most similar with the very late colonising CF strains (average mash distance between PAO1 and Pa422, Pa423, Pa423.1, Pa427, Pa428, Pa436 was 0.048, Table S7), suggesting this could be a good proxy of the response of the strains isolated at later stages in the infection. We observed that PAO1 caused a 55.8% reduction in the biomass weight compared to the negative control and only 2 out of 9 germinated seeds. When CF strains were inoculated, root and shoot development was partially or completely abolished and seed/seedling weight was reduced when infected with most of the clinical *P. aeruginosa* strains (109 CF strains caused an average weight reduction of 54.2% compared to negative controls and an average germination rate of 1.9 seeds out of 9; Figure S1). However, we also found 10 strains that reduced the seed weight less than 33.3% (Pa76, Pa87, Pa205, Pa206, Pa214, Pa223, Pa389, Pa243, Pa412.1 and Pa440), indicative of reduced virulence relative to PAO1 (Figure S1).

The 119 strains were then classified into early, intermediate, late and very late adapted strains based on the exact timepoint in the infection history of the individual CF patient each of these strains was isolated from (see material and methods). It was observed that clinical strains that had colonised CF lungs for the longest time showed relatively higher virulence in mung beans compared to early isolates (R^2^=-0.3, p-value=9.5E-04 for weight, Figure 4A). These results suggest that mung bean model was good at predicting the virulence of chronic, lung-adapted CF strains.

**Figure 4.**
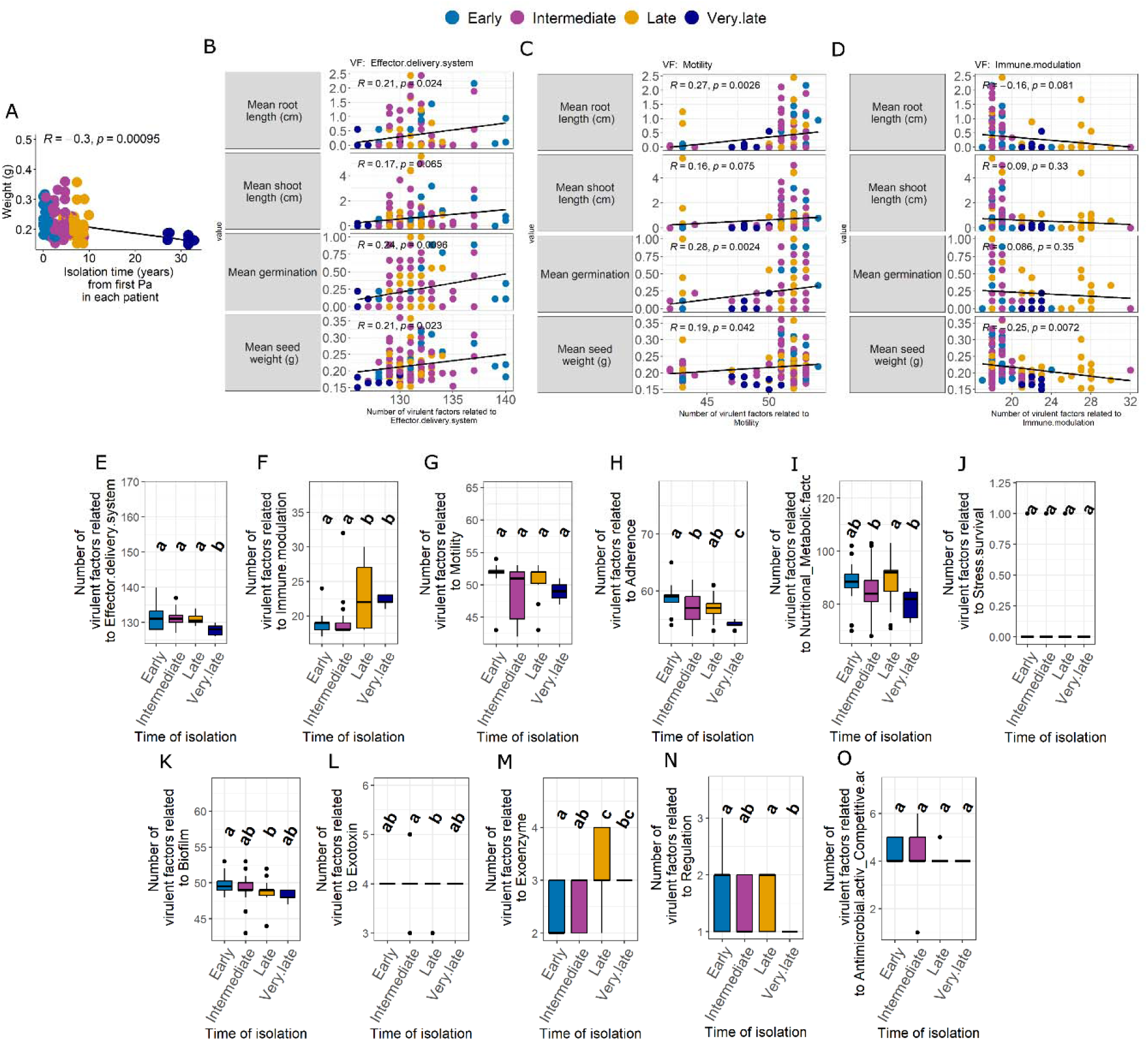
Virulence of 119 clinical *P. aeruginosa* strains in mung beans and correlation with virulent factors/genes and time of isolation. **A.** Pearson correlation between the seed weight and the isolation time of the 119 strains, **C-E**. The graphs report Pearson correlations between the number of effector delivery genes **(C**), the number of genes involved in motility (**D**) and the number of immune modulator genes (**E**) and the four virulence traits measured in mung beans. Traits were root and shoot lengths in cm; germination with 0 non-germinated seeds and 1 germinated seeds and weight of the seeds/seedlings at 5 dpi. Dots represent the average of that measurement in 9 mung bean seeds for each strain. Higher values indicate lower virulence, values closer to 0 indicate higher virulence for all traits. **F-P.** Number of genes involved in effector delivery (**F**), immune modulation (**G**), motility (**H**), adherence (**I**), nutritional and metabolic factors (**J**), stress survival (**K**), biofilm (**L**),exotoxin (**M**), exoenzyme (**N**), regulation (**O**) and antimicrobial activity and competitive activity (**P**) in the 119 strains classified according to the isolation time from each patient into early (or acute), intermediate, late or very late stages (representing more chronic stages). Different letters indicate statistically significant differences according to a post-hoc Tukey’s Honest Significant Difference test performed after ANOVA analysis.

To explore which genes could be associated with mung bean virulence, we first constructed a *P. aeruginosa* pangenome with Roary using PAO1 as a reference genome, creating a matrix of presence/absence of orthologus genes in all 119 strains. The pangenome was annotated using Prokka and the annotations were blasted against the full Virulence factor database (https://www.mgc.ac.cn/VFs/) to identify potential virulence genes. The matches were divided into the categories reported by the database: Adherence", "Antimicrobial activity/Competitive advantage", "Biofilm", "Effector delivery system”, “Exoenzyme", "Exotoxin", "Immune modulation”, "Motility", “Nutritional/Metabolic factor", "Regulation” or “Stress survival". We then performed a Pearson correlations analysis between the number of virulence genes (per category) and virulence measured in mung beans using all virulence-associated traits (root and shoot length, germination and weight of the seeds) were then calculated. We observed that reduction in virulence was associated with an increase in the number of effector delivery systems and motility genes (R^2^=0.21, p-value=0.024 for root length, R^2^=0.17, p-value>0.05 for shoot length, R^2^=0.24, p-value=0.0096 for germination rates, R^2^=0.21, p-value=0.023 for seed weight, Figure 4B and R^2^=0.27, p-value=0.0026 for root length, R^2^=0.16, p-value>0.05 for shoot length, R^2^=0.28, p-value=0.0024 for germination rates, R^2^=0.19, p-value=0.042 for seed weight, Figure 4C). Moreover, we observed that increase in virulence (lower weight of the seeds) was correlated with a higher number of genes related to immune modulation (R^2^=-0.25, p-value=0.0072, Figure 4D). Especially, the strains isolated at very late stages of the CF infection carried the lowest number of effector delivery genes (F(_3,115_)= 5.58, p-value=0.001; Figure 4E), while late and very late strains carried the highest number of immune modulation-related genes (F(_3,115_)= 18.54, p-value=7.06e-10; Figure 4F). For example, *wbpA* gene (pangenome_7205) associated with B-band LPS biosynthesis(74) was only present in very late strains (Figure S2)[85]. Despite no significant changes being observed for motility genes (Figure 4G), fewer number of genes linked with other virulence factors such as adherence, metabolism, biofilm or regulation were carried by strains isolated at late and very late stages (Figure 4H-O). Overall, these results suggest that CF strains isolated at later stages of infections were relatively more virulent in the mung bean, and that these strains carried a fewer number of genes associated with effectors, adherence, metabolism, biofilm, or regulation genes but a higher number of genes associated with immune modulation.

## DISCUSSION

In this work, we demonstrate that the infection of *P. aeruginosa* in mung beans caused expression of several virulence-related genes following different steps of infection, including PAO1 motility, chemotaxis, phosphate regulation and biosynthesis of pigments. Concomitantly, mung beans upregulated genes involved in defense responses and downregulated genes involved in photosynthesis, suggesting allocation of resources towards the immune system. Moreover, some expressed genes were specific to the mung bean model and included Type II secretion system and phosphate starvation genes, which might be required for successful plant infections. PAO1 core-transcriptomic gene expression was more similar to ex-vivo sputum, artificial media, burns or excisions in mouse and rabbit, but less similar to PAO1 gene expression changes in human bronchial cells, mouse lungs, or SCFM media. We also confirmed that mung beans could be used as a screening method to distinguish virulence differences between *P. aeruginosa* CF strains, where strains isolated from later stages of CF infections showed higher levels of virulence and were associated with higher number of genes linked with host immune modulation. Together these results suggest that the mung bean model could potentially be used to quantify the virulence of chronic versus early CF strains.

We first characterised the transcriptomic response of PAO1 during the mung bean infection. We observed that PAO1 gene expression was sequential including increases in gene expression abundance at later time points for some genes, compared to early time points. For example, *wbpE*, which is involved in B-band O-antigen biosynthesis (48, 49), and *cheY*, a regulator of chemotaxis (51), showed higher expression at 30hpi and 48hpi. PAO1 chemotaxis-associated genes were also induced when infecting poplar trees (44) At 48hpi, we also observed induction of several genes involved in phosphate import such as *phoU*, part of the phosphate regulon*, pstS,* part of an ABC transporter system, and the porine *oprO*. Phosphate uptake-related genes such as the porins oprO and oprP have been reported to change their expression levels distinctly when PAO1 infected susceptible compared to resistant plants (41). The induction of phosphate starvation-associated gens can imply a shift to virulence phenotype (53, 54). At 48hpi, we also observed an induction of genes associated to quorum sensing, such as the gene PhzA/B-like protein involved in phenazine biosynthesis that acts as a signalling molecule (55) activating the quorum-sensing genes (77). Finally, at 48hpi and 72hpi, the gene *clpP2*, a protease, involved in alginate production and biofilm formation (57, 58) showed increased expression. Overall, the PAO1 response in mung bean matched Harrington et al. (2025) (78) results, who reported a significant increase in the expression of phenazine biosynthesis and quorum sensing between 48hpi and 72hpi in a CF lung model in pigs when inoculated with the *Pseudomonas aeruginosa* PA14 type strain. PAO1 gene expression responses in mung bean also resembled results by D’Arpa et al. (2021), who reported the gene expression changes of PAO1 in skin wounds. D’Arpa et al. (2021) showed colonisation-related genes were activated at early hours after infection (2h and 6h), whilst polysaccharide biosynthesis, responses to stresses and early biofilm formation happened at 24hpi (79, 80). On the other hand, the plant response at the early stages after infection was characterised by induction of immune responses and repression of growth and development- a common effect when combating a pathogen (81).

We then used comparative analysis as reported in (82) to find similarities and differences between gene expression changes in mung beans and other virulence models including *in vitro* and *in vivo* samples. The 24 samples obtained from 4 previously published studies (63–66), were further categorised by model type: *in vitro* media, media with epithelial cells, lung cells from human and animal, sputum (artificial and ex-vivo), models that require burns or excisions (classified as skin models), or mung beans. We identified 1172 genes, 20% of the total PAO1 gene content, that were expressed in all the 6 models and were enriched in functions that were associated with general maintenance and growth of the cells, such as transcription or cell division, but also stress responses. Importantly, these core-transcriptional genes were not expressed in the same way in all the conditions, but the expression was more similar between mung bean, skin-models, *in vitro* media and sputum models than lung cell models. This highlights that PAO1 gene expression responses in mung bean are distinct from other models, probably due to differences in immune systems, cell types and biochemical environments between human, animal and plants. We also observed 202 PAO1 genes, which were only expressed in mung bean model. These included several genes of the *arn* operon, associated to resistance to the antibiotics polymyxin and colistin, due to the modification of the lipopolysaccharide with 4-amino-4-deoxy-L-arabinose, reducing the negative charge of the bacterial membrane (67). We also found genes involved in the transport of microcins, or low-molecular-weight peptide molecules with antibacterial activity (69); Type II secretion systems (gsp operon), crucial to export enzymes into the plant during infection (83, 84), and phosphatase *phoA*. These findings highlight that PAO1 uses specific virulence genes to establish infection when colonising plants versus animals. Overall, mung beans were proven to be a good model for assessing *P. aeruginosa* infections; however, further studies should focus on shedding light to all the infection mechanisms used by *P. aeruginosa* in plants and whether other plant species could be considered as virulence models.

We finally tested if the mung bean model could differentiate virulence variation between 119 clinical CF strains and compared to PAO1. As PAO1 was found to be the most virulent strain in mung beans and showed the most similarity to strains from later stages in terms of mash or genetic distance, our transcriptional study could become a proxy for assessing the virulence of CF strains. We also found that the strains isolated from later stages of the CF infection showed higher virulence in mung beans than the strains isolated from early stages of CF infections. These results seemingly contrast with studies reporting loss of virulence of chronic strains in other virulence models (85–87), due to loss of virulence factors or accumulation of mutations in those genes seen in chronic CF infection strains compared to acute strains. To link virulence variation with the presence of different virulence genes, we compared the genomes of the 119 strains and identified the number of virulence genes in each strain. We observed that the number of virulence genes, including effector delivery systems, motility, biofilm, regulatory elements, and genes involved in stress or adhesion, was significantly lower in strains isolated at later stages of CF infection, in line with previous studies. However, we also observed that the most virulent strains (late and very late strains) had an increased number of virulence factors related to immune modulation, including genes involved in LPS biosynthesis, such as *wbpA*, the first enzyme in the biosynthetic route of 2,3-diacetamido-2,3-dideoxy-D-mannuronic acid, which is an essential part of the B-band O-antigen of *P. aeruginosa* PAO1 (74). The absence of this gene in the early and intermediate strains might imply that the B-band is absent. These results suggest that late and very late strains had undergone pathoadaptations, resulting in loss of some virulence factors, but that the relatively higher number of immune modulator genes could be associated with increased virulence of chronic strains in the mung bean model. These adaptations could also help the chronic strains to combat the host immune system, explaining why they can persist in chronic infections. Importantly, despite suffering pathoadaptations, the higher virulence of late and very late strains could also be associated with distinct gene expression of the virulence factors and immune modulators in mung beans compared to other models.

To conclude, we reported the first PAO1 transcriptomic response in mung beans and showed specific gene expression changes associated with the infection process. Our results confirm that mung beans could be used as a quick screening method to assess *P. aeruginosa* virulence, using straightforward measurements that include checking germination, the weight of the seed at 5 dpi and measuring stem and root length, and substituting inoculation in models that require live animals, reducing ethical considerations and speeding up the results. Mung bean was especially good at identifying chronically virulent CF strains that had gone through significant pathoadaptations in the CF lung. The mung bean model could hence offer an additional virulence model to screen bacterial pathoadaptations and provide novel insights on *P. aeruginosa* pathogenicity in plants.

## MATERIAL AND METHODS

### Bacterial strains

We selected *P. aeruginosa* PAO1 strain, which was kindly donated by Marvin Whiteley, and 119 *P. aeruginosa* isolates provided by Helle Krogh Johansen from a longitudinal cystic fibrosis airway study (12) for our study. The CF strains, named by their sequence ID corresponding to previously published data (‘Pa’ for P. aeruginosa has been added to the names for easier identification)(88) were classified into different lineages based on their genetic similarity as described previously (12) and included strains from 9 patients. Strains from patient 2, patient 6, patient 8 and patient 9 were sampled from 0 years post infection to 4.1, 7.5, 7.3, and 6.7 years, respectively, and had an average length of infection of 2.4, 5.1, 3.8 and 3.2 years. Strains from the patient 1, patient 5 and patient 3 were sampled at 0.2, 0.4 and 1.4 years to 6.7, 5.2 and 8.9 years post-first infection with an average of 3.3, 3.5 and 5.7 years, respectively. Finally, the strains isolated from the patient 7 and patient 4 were sampled from years 4.4 and 27.1 to 9.8 to 32.8 years post-first infection and showed an average of 7.5 and 29.9 years, respectively. Moreover, strains were classified into early, intermediate and late infection strains based on the length they had been infecting individual patients, which was achieved by dividing the length of the CF infection within each patient into three equally long time periods (Table S1). As even the first strains from patients P36F2 and P82M3 were isolated several years after the typical time of the first infection, and the average time of sampling was longer than in the other patients, we classified all of them as “very late” adapted strains.

### Mung bean infections

Mung beans (*Vigna radiata*) were obtained from Moles Seeds UK Ltd (Colchester, United Kingdom). The seeds were disinfected by submerging them in ethanol 70% for 2 minutes, followed by rinsing in sterile water. To start germination, the seeds were placed on water-agar (1%) plates and maintained at room temperature for 24h. Bacterial cultures were prepared by inoculating 5 mL of LB with streaks from strain glycerol stocks kept at -80°C and grown at 37°C for 24h with agitation of 100rpm. Cultures were then centrifuged at 5000*g* for 5 minutes and the pellet was resuspended in 5mL PBS. Colony counts were calculated from this suspension using serial dilutions followed by plating and the initial bacterial concentrations were adjusted to approximately 1.3E+10 CFU/mL. To infect the mung beans, the seeds were submerged in bacterial PBS-cultures for 24h at 30°C in static conditions. Negative control mung bean seeds were submerged in 5 mL of sterile PBS without bacteria in otherwise similar conditions. After infection, seeds were surface sterilised with ethanol, rinsed with water and transferred to a 15mL Falcon tube containing 5mL of soft-agar (7.5g agar/L) Murashige Skoog Basal Medium (MS). The tubes were then moved to a PHCbi growth cabinet (MLR-352) and kept atÕ28°C with 16h light/8 dark cycle for a maximum of 5 days. After finishing the experiment, the plants were heat-treated for 5 minutes at 60°C to detach them from the soft agar, allowing measuring changes in the root and stem lengths (cm), germination rates (1 when germinated and 0 when no germination happened) and weight (g). Changes in the loss of biomass were plotted using ggplot2 in R. Differences between strains were compared using the non-parametric Kruskal-Wallis test, followed by the Dunn test from the FSA in R (89). The mean of each of the 9 replicates of each of the 4 traits (scaled values) was calculated for each *P. aeruginosa* isolate.

### Assessing *P. aeruginosa* PAO1 and mung bean gene expression changes during early infection

To identify *P. aeruginosa* gene expression during the early stages of mung bean infection, we focused on PAO1 strain, due to its high virulence in mung bean and the high number of publicly available PAO1 transcriptomic datasets that allow us to compare gene expression difference between PAO1 in mung bean and other models (see below). For each time point sampled, we used 3 infected and 3 uninfected seeds as replicates, which were destructively sampled from a larger set. The 24h infection of control matched with the time before moving the seeds to the soft agar MS media, and further destructive sampling was conducted at 30h, 48h and 72h time points. The RNA (3 biological replicates at each time point for infected and uninfected control samples) was extracted by first grinding the individual seeds in liquid nitrogen with a mortar and pestle and using the EZNA Plant RNA kit for RNA extractions (Omega Bio-tek, Georgia, USA) following the manufacturer’s instructions. The RNA quality was quantified using the Agilent Technology 2,100 Bioanalyzer (Agilent Technologies, CA, USA) and the total RNA was used for metatranscriptomics analysis of both bacteria and plant. Sequencing was performed with Illumina by the external services of Novogene UK using a NovaSeq X Plus Series instrument (PE150). One of the biological replicates of the infected samples at 24h (one seed) could not be sequenced due to low RNA concentration and was removed from the analysis. After sequencing, all the reads were trimmed using trimmomatic 0.39 (90). For the bacterial analysis, we mapped reads back to PAO1 reference genome with salmon (91) using minScoreFraction 0.9 to increase the mapping rates.

To visualise global gene expression changes, Principal Component Analyses (PCA) were used with the function prcomp of the stats package in R on the log(TPM+1). For the bacterial reads, a weighted gene co-expression network was constructed using the package WGCNA(92) in R with the transcript per million (TPMs) of the 5748 genes after removing genes with missing values using goodSamplesGenes function. The power was set to 20 to fulfil scale-free topology requirements, as only 11 samples were used to build up the network. The parameters TOMType = "unsigned", minModuleSize = 20,networkType = "unsigned", reassignThreshold = 0, mergeCutHeight = 0.25,deepSplit = 4 were used to create the network using the function blockwiseModules. Gene ontology enrichment was assessed using topGO (93) package in R to identify functions associated with virulence in each module. Only 6 modules were further selected according to their association with virulence-related functions. The networks of these modules were exported to Cytoscape (94) using exportNetworkToCytoscape function.

The Cytoscape function “Analyse network” was used to calculate the degree or the number of connections of each gene in these 6 modules. Only the genes with the highest degree in each of the selected modules were considered as hub genes with potential regulatory functions during the infections and were further explored. Heatmap of the expression log (TPMs+1) were obtained using Heatmap from the package ComplexHeatmap in R. Statistical differences between the different time points were assessed using ANOVA using the function aov from the package stats in R, followed by the Tukey’s Honestly Significant Difference (HSD) test.

For the mung bean transcriptomic analysis, we mapped the trimmed reads against the mung bean genome Vrad_JL7 (95) CDS models with default parameters. Then, a PCA was performed as above to assess global transcriptional changes between infected and non-infected samples. We further identified differentially expressed genes using DESeq2 (96) with the lfcShrink function and method apeglm (97), where genes were considered to be upregulated when log2FoldChange ≥ 2 and padj <= 0.05 and downregulated when log2FoldChange ≤ - 2 and padj <= 0.05. Samples from 24h were not included in the analysis due to only having 2 replicates in the bacterial treatment samples. Gene ontology enrichment, using the annotation provided by (95), was assessed using topGO package in R (93). We further selected the 5 most upregulated and downregulated as well as the 5 genes with lowest p-adjusted values for each of the 3 contrasts (30hpi, 48 hpi and 72hpi) and differences in the expression of selected genes were assessed as above using ANOVA and HSD test.

### Quantification of PAO1 densities during mung bean infections using qPCR

To quantify *P. aeruginosa* densities during mung bean infection, we used the same RNA extracted at 24 hpi, 30 hpi, 48 hpi and 72 hpi (3µL/sample) to obtain cDNA using SuperScript™ IV Reverse Transcriptase (ThermoFisher Scientific), with RNaseOUT and random primers following the manufacturer’s instructions. The cDNA amount was quantified with a NanoDrop™ 2000 (ThermoFisher Scientific) and all the samples were diluted to 100 ng/µL. In parallel, PAO1 was grown for 24h in LB as above and the harvested cells were used to extract the DNA using the E.Z.N.A.® Bacterial DNA Kit according to the manufacturer’s instructions. PAO1 DNA was used to create a standard curve ranging from 7.6 ng/µL to 75.6 fg/µL. qPCR was performed for mung bean samples using iTaq Universal SYBR Green Supermix and a final concentration of 20 µM of the primers set PA14 forward 5’-CGGTACAGGTCGGCACG-3’ and reverse 5’-CGAGGGACGAAGGTAAGGA-3; designed by (98) and amplifying a hypothetical protein. Each mung bean sample was quantified in technical triplicate, and the average of the three technical replicates was used as the Ct for each sample. PAO1 DNA concentrations were calculated using the standard curve: *log(DNA concentration)=-3.5669 Ct +18.768*; R^2^ =0.9961.

### Comparative transcriptomics to identify *P. aeruginosa* genes that are expressed similarly in different virulence models

To assess if the responses of PAO1 in mung bean were similar to other commonly used *in vitro* and *in vivo* virulence models, we compared our data with paired-end and single-read RNA-Seq experimental data from previously published studies (Table S2, (63–66)). These data sets and our mung bean transcriptome, were categorised into 13 different submodels within 6 types of virulence models: 1) “*In vitro* media” (this model included artificial sputum media, *Pseudomonas* Isolation Agar (PIA), M63, MOPS or synthetic CF sputum medium (SCFM); 5 samples, each with multiple replicates and a total of 17 samples), 2) “media and lung cells” (this model included samples with SCFM2 and human bronchial epithelial cells named epiSCFM.human; 3 samples with 3 technical replicates in each), 3) “Lungs” (including human bronchial cells and mouse lungs; 2 samples with 3 technical replicates each), 4) “Sputum” (ex-vivo sputum 6 patients with 1 technical replicate each, all grouped in the analysis), 5) “Skin” (including murine burn wounds (1 sample with 2 technical replicates), murine chronic wounds (1 sample with 2 technical replicates) and rabbit wounds (4 samples corresponding to a time series and two technical replicates in each)) and 6) “Mung bean”. PAO1 transcriptomic responses in these models were compared with PAO1 responses in the mung bean samples collected at different time points (“Mung beans”, 11 samples at 4 time points). For the mung bean experiment, the reads were trimmed and mapped against the PAO1 reference genome as explained earlier, keeping minScoreFraction set to 0.9. The mapping (Table S2) ranged from 0.03% to 98.92%, with an average of 35.34%; however, as some of the mung bean time points (24h) also showed low mapping rates likely due to low bacterial growth (minimum of 0.14%), we kept all the samples in the analysis. A PCA was conducted to observe differences in the gene expression between different samples, submodels and types of virulent models, by first obtaining log(TPMs+1), followed by the function normalize.quantiles of the packages preprocessCore, followed by PCA using the prcomp function of the stats package in R. perMANOVA was also calculated using pairwise.adonis() from the package pairwiseAdonis in R (99), using the same quantile and log normalised data, with Bray-Curtis distance, 999 permutations to confirm differences across model types, considering the expression of the 5765 genes.

To identify genes that were expressed in the 6 different types of virulence models, we first calculated the TPMs of all samples within virulence models. Only genes expressed at least in 1 TPM in a sample, were considered expressed in that sample. A gene was considered expressed in a submodel when it was expressed in at least 75% of the samples within the submodel. In a similar way, a gene was considered expressed in a model type if it was expressed in 75% of the samples within that model type. These filters identified 4190 genes expressed in media, 5368 in media and lung cells, 3244 in lungs, 5136 in sputum, 1581 in skin and 5381 in mung bean. UpSet package in R was used to identify genes that were expressed in all 6 types of virulence models, resulting in total of 1172 genes expressed across all virulence models. To avoid biases due to the use of different sequencing technologies and platforms, we normalised the expression of these 1172 genes using quantile normalisation of the log (TPM+1) with the function normalize.quantiles from the package preprocessCore in R and plotted in a heatmap using the function heatmap in R. We then used the average of the normalised expression of the 1172 genes to find which treatments were more similar in their gene expression by using a Pearson correlation with the function cor from package psych, transforming the correlation to distance using the function as.dist. Samples were then clustered according to this distance by using hclust function and average method and visualised using a heatmap. In addition, gene ontology enrichment analysis was performed as earlier described to identify the functions of 1172 genes that were expressed in all virulence models, and gene ontology enrichment was also performed separately for 202 genes that were only expressed in the mung bean model (identified based on UpSet package in R).

### Assessing the virulence of clinical *P. aeruginosa* CF isolates using mung bean model

Previously (46) reported that inoculating *P. aeruginosa* will reduce the weight and growth of mung bean seeds. To assess how well mung beans can distinguish virulence differences between clinical *P. aeruginosa* strains, we tested the virulence of 119 *P. aeruginosa* CF strains individually. The mung bean inoculations were performed as described earlier with the seeds maintained for a total of 5 days in the plant growth cabinet (N=9 mung bean/strain; negative control seeds without bacteria were kept in sterile media, N=27). Four virulence traits were measured as explained above: root and shoot length, germination rates (1 when the seed germinated and 0 when it did not) and weight of the seeds at 5 DPI.

### The DNA extraction and genome assemblies of clinical *P. aeruginosa* CF isolates

To link genomic variation in CF strain virulence genes with virulence measured in the mung bean model, PAO1 and twenty-three CF strains (Table S3) were selected for long-read Nanopore sequencing as they belonged to 23 different lineages based on a previous study (12). The reads of 5 of these strains were already used to assess prophage diversity in the genome . Please note the PAO1 genome corresponds with a PAO1 strain provided by Marvin Whiteley (100). The CF strains originated from 9 patients and were collected longitudinally across a period of infection of a maximum 88 months (Table S4). Bacteria were grown in LB from glycerol stocks stored at -80 °C maintained at 37 °C and agitation of 100rpm for 24h. Cultures were centrifuged at 5000 *g* for 5 minutes to collect cells. Genomic DNA was isolated using the Qiagen Genomic tip 100/G kit following the manufacturer’s instructions (Qiagen, Manchester, UK). Sequencing was performed either by the University of York’s Technology Facility using Oxford Nanopore Minion Flowcell (R9.4.1) or at the sequencing facility of the NNF Center for Biosustainability (now renamed BRIGHT), Technical University of Denmark. Raw reads were demultiplexed and trimmed. Base calling was performed using Guppy v6.3.9 with the super accuracy model. Reads were then assembled with Flye v2.9 using a genome size parameter of 7 million, followed by three rounds of medaka v1.9.1. Finally, Illumina reads of the same isolates obtained by (12)(BioProject PRJEB5438) were trimmed using Trimmomatic v0.39 and genomes were polished using pilon v1.23. Seqkit v2.3.1 was used to calculate genome statistics and BUSCO v.5.4.3 to assess genome completeness against the bacterial database. The genomes of sequenced *P. aeruginosa* strains were assembled into 1 contig, except for Pa382, which was assembled into 4 contigs, Pa182, Pa199 and Pa339, which were assembled into 3 contigs and Pa200 and Pa390, which were assembled into 2 contigs. The smallest contig size in those strains was 14,685 bp, suggesting the presence of plasmids. The minimum genome size was 6,195,234 bp and the maximum size was 7,170,410bp, with GC% between 65.72% to 66.61% (Table S3) with a completeness ranging from 99.2% to 100% (Table S3). Prokka v.1.14.5 was then used for *de-novo* annotation of each genome.

The remaining previously sequenced 96 CF isolate genomes were re-assembled by scaffolding the publicly available Illumina-based contigs (12) with the lineage-specific complete genomes assembled in this study using SibeliaZ v1.2.6 followed by Ragout v2.3, to reorder the contigs into scaffolds. Dnadiff was used to align the old assemblies with the new scaffolded genomes and the number of aligned and unaligned base pairs and contigs were calculated. We obtained an average of 64,956 bp of unaligned base pairs of the new genomes, which corresponds only to 0.98% of the new genomes’ length. This suggests the scaffolding allowed more contiguous genomes without loss of genetic information (Table S5-S6). Mash distance was used and calculated as a measure of genetic distance between all 120 strains (119 CF strains and PAO1; Table S7) using MASH sketch and triangle functions (101).

### Identification of virulence genes associated with mung bean virulence in clinical *P. aeruginosa* cystic fibrosis isolates

To identify virulence genes in newly assembled CF isolates, the genomes of the 119 strains and PAO1 were first input into Roary v3.13 to create a matrix of presence and absence of genes, after Prokka v.1.14.5 was used to annotate the reference pangenome. We then used Blastn to match the pangenome annotations with the DNA full virulence factor database (https://www.mgc.ac.cn/VFs/). To obtain stringent results, only Blast matches with an E-value<0.01, a length of 100 bp and an identity of more than 70% with the database were kept for further analysis. All blastn matches were further divided into one of the following categories based on the virulence factor database: Adherence", "Antimicrobial activity/Competitive advantage", "Biofilm", "Effector delivery system”, “Exoenzyme", "Exotoxin", "Immune modulation”, "Motility", “Nutritional/Metabolic factor", "Regulation” or “Stress survival". Finally, to test if some of these genes were associated with virulence in mung bean, we performed a Pearson correlation using the stat_cor function in R between the number of genes in each of the virulence categories and the observed virulence measured in mung bean model (root length, stem length, germination and weight of the seeds).

### Statistical analysis

Statistical differences between groups were assessed in R 4.3.3, using ANOVA or Kruskal-Wallis test according to the normality of the data, followed by Tukey’s HSD test from the rstatix package or Dunn’s post hoc test from the package FSA with Bonferroni correction for multiple comparisons.

## ACKNOWLEDGMENTS

This project was undertaken on the Viking2 Cluster, which is a high-performance computer facility provided by the University of York. We are grateful for computational support from the University of York High Performance Computing service, Viking and the Research Computing team. The authors would like to acknowledge the contribution of Dr. Sally James and Dr. Lesley Gilbert from the Bioscience Technology Facility at the University of York for their assistance in the sequencing. We kindly thank Marvin Whiteley for providing the PAO1 strain used in this study.

## AUTHOR CONTRIBUTIONS

SFO performed all the plant experiments, extracted RNA, performed all the bioinformatic analysis and wrote the first draft. EH ad IK sequenced the genomes of the strains. SFO, JWBM and VPF conceived and planned the project. CSM and VPF obtained funding for the project. SFO, JWBM and VPF wrote the manuscript, and all authors reviewed it.

## DATA AVAILABILITY

The Oxford Nanopore raw data for the genomes is available in NCBI Sequence Read Archive (SRA) BioProject PRJNA1347034, whilst the raw data of the metatranscriptomics is available in PRJNA1359585. All the scripts can be found at https://github.com/sfortega/Pseudomonas_pathoadaptations.

## ETHICS

The use of clinical *P. aeruginosa* isolates was approved by the Capital Region’s Committee on Health Research Ethics (H-21078844).

## SUPPLEMENTARY FIGURES

**Figure S1.**
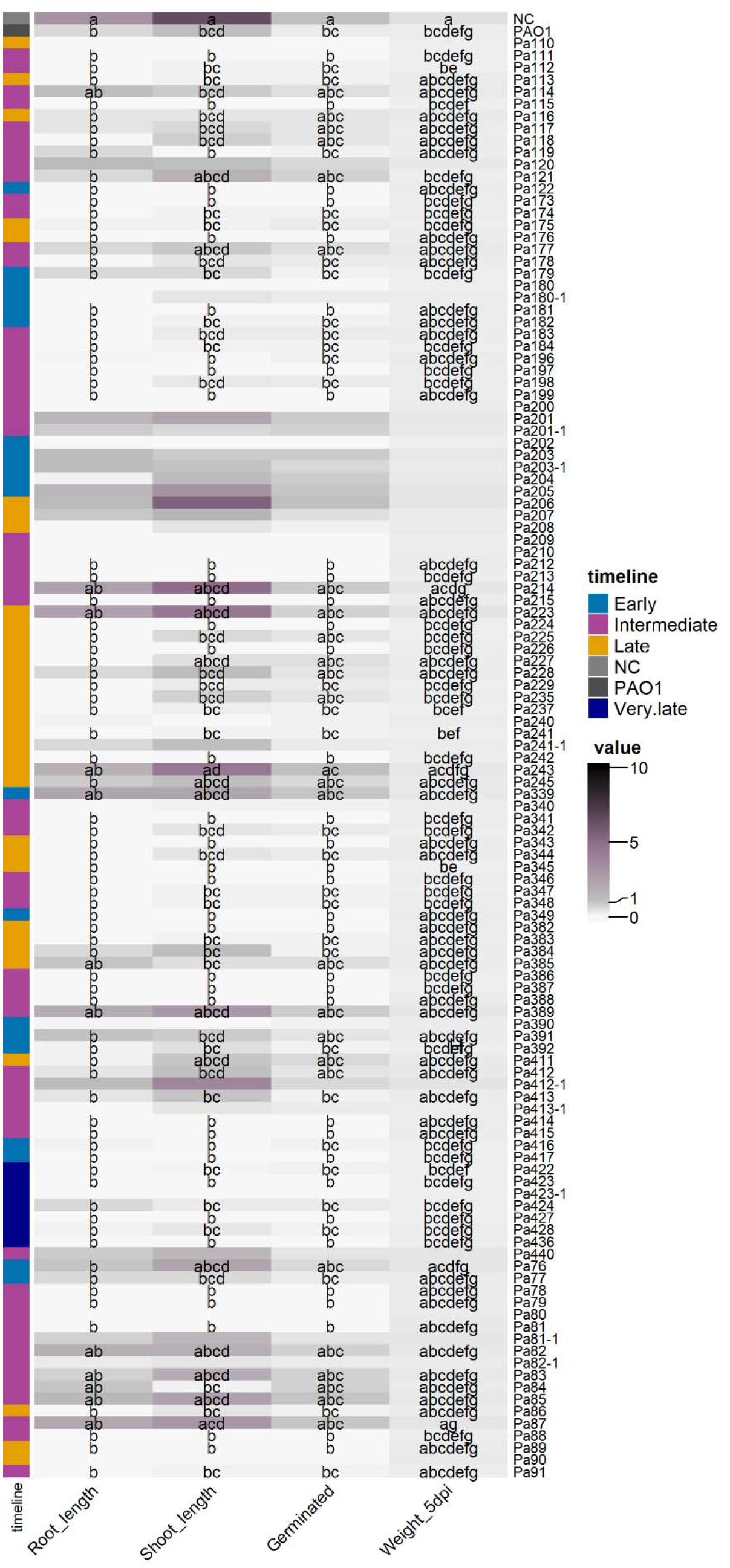
Changes in root and shoot length in cm, germination (being 0 not germinated and 1 germinated) and weight measured at 5 dpi (in g) relative to negative control and PAO1 treatments. Different letters indicate statistically significant differences according to a post-hoc Dunn’s test performed after Kruskal–Wallis analysis. **B**

**Figure S2.**
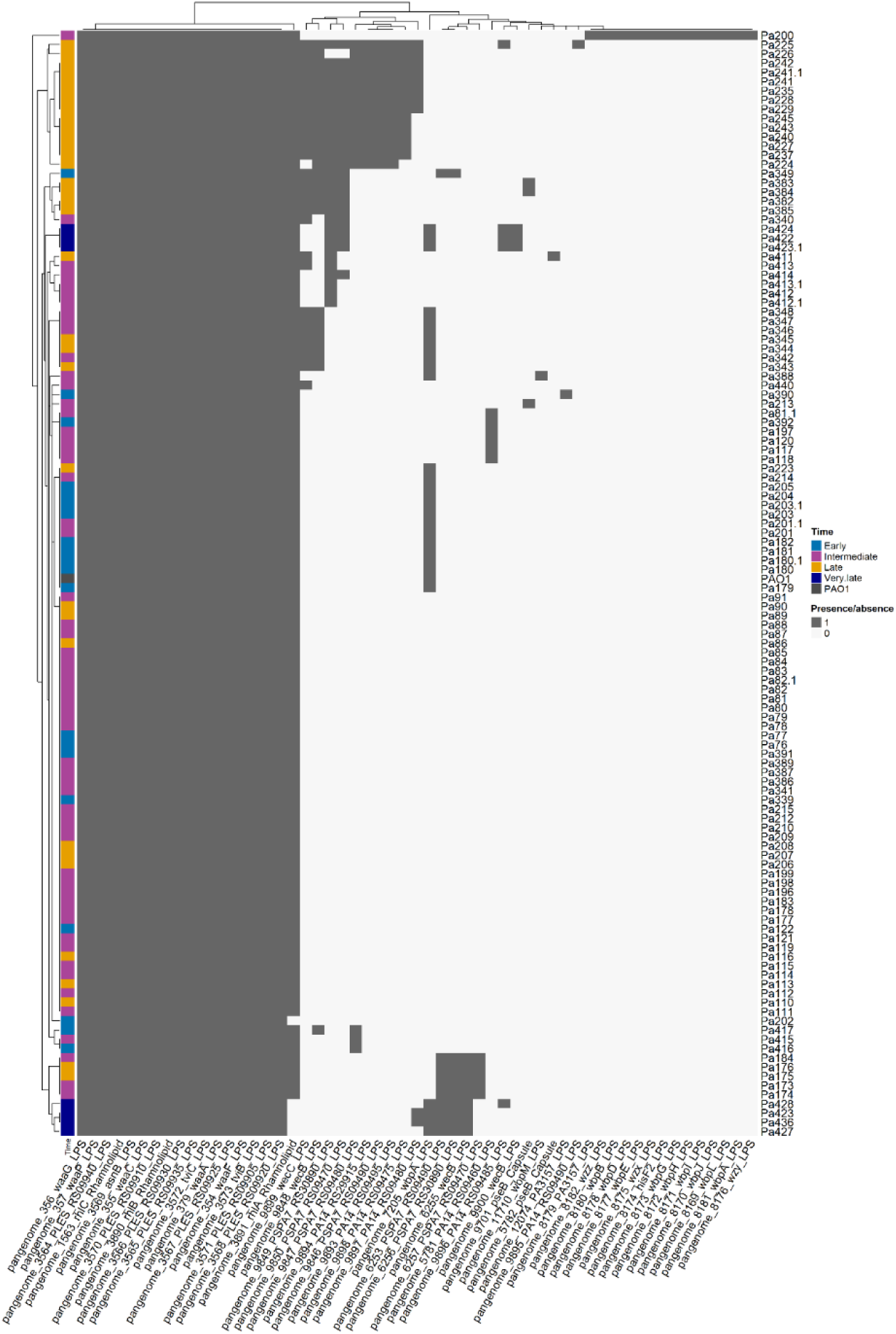
Heatmap of the presence and absence of virulence factors identified as Immune modulators in the 119 CF strains and PAO1.

## SUPPLEMENTARY TABLES

Table S1. P. aeruginosa clinical strains. Sequence ID corresponds with previous published data, but Pa has been added to the names for easier identification.

Table S2. RNA-Seq studies representing the response of PAO1 in different virulence models.

Table S3. Genome stats of 23 P. aeruginosa strains assembled with ONT and Illumina data.

Table S4. Strains according to the timepoint of sampling in each patient

Table S5. Genome stats of 96 P. aeruginosa strains assembled using a reference in Table S3.

Table S6. Comparison between the previously assembled genomes and the new genomes for the 119 clinical P. aeruginosa strains.

Table S7. Mash distance between the 119 CF strains and PAO1.

Table S8. PAO1 DNA amount in the mung bean seeds at early time points after infection (24 hpi, 30 hpi, 48 hpi, 72 hpi)

Table S9. Mapping percentages to PAO1 and Mung bean JL7.

Table S10. WGCNA results created with the PAO1 genes, reporting gene significance (GS) and p-value of each gene at the different time points after infection (24 hpi, 30 hpi, 48 hpi, and 72 hpi) and gene annotations.

Table S11. Gene ontology enrichment (only biological processes) of some modules in the WGCNA assessing PAO1 gene expression over time.

Table S12. Degree of each gene in the black module of the gene co-expression network of the different time points after inoculation of PAO1 in mung beans.

Table S13. Degree of each gene in the blue module of the gene co-expression network of the different time points after inoculation of PAO1 in mung beans.

Table S14. Degree of each gene in the brown module of the gene co-expression network of the different time points after inoculation of PAO1 in mung beans.

Table S15. Degree of each gene in the turquoise module of the gene co-expression network of the different time points after inoculation of PAO1 in mung beans.

Table S16. Degree of each gene in the salmon module of the gene co-expression network of the different time points after inoculation of PAO1 in mung beans.

Table S17. Degree of each gene in the tan module of the gene co-expression network of the different time points after inoculation of PAO1 in mung beans.

Table S18. Differentially expressed genes analysis with upregulated genes in blue and downregulated genes in red at 30 hpi.

Table S19. Differentially expressed genes analysis with upregulated genes in blue and downregulated genes in red at 48 hpi.

Table S20. Differentially expressed genes analysis with upregulated genes in blue and downregulated genes in red at 72 hpi.

Table S21. Gene ontology enrichment analysis of the upregulated and downregulated genes in mung beans after infection with PAO1.

Table S22. TPMs of the different virulent models.

Table S23. Gene ontology enrichment analysis of the 1172 genes expressed in all 6 virulence models and the 202 genes only expressed in mung beans.

Table S24. Virulence of the 119 P. aeruginosa strains in mung beans, reporting root length, stem length and weight at 5 days post-infection.

